# Overcoming migratory exhaustion: Expression of Matrix metalloproteinase-8 (MMP-8) promotes tumor infiltration and anti-tumor activity of CAR-T cells

**DOI:** 10.1101/2023.06.21.545815

**Authors:** Irfan N. Bandey, Melisa J. Montalvo, Harjeet Singh, Navin Varadarajan

**Affiliations:** Department of Chemical and Biomolecular Engineering, University of Houston, Houston, Texas 77204, USA; Division of Pediatrics, University of Texas MD Anderson Cancer Center, Houston, Texas 77030, USA

## Abstract

Despite the encouraging success of chimeric antigen receptor (CAR) T-cell therapy in treating hematological malignancies, the translation of adoptive cell therapies to solid tumors remains a challenge. Several studies have attributed the inability of tumor-infiltrating T cells to traffic to solid tumors, primarily to the presence of the extracellular matrix (ECM) and immunosuppressive environment of solid tumors. The ability of the transferred T cells to infiltrate the tumor is an essential prerequisite for anti-tumor activity. We show here that upon activation and expansion, T cells quickly lose their migratory capacity, leading to migratory exhaustion. At the molecular level, migratory exhaustion could be attributed to the downregulation of matrix metalloproteinase 8 (MMP8). To overcome this, we hypothesized that T cells genetically modified to secrete the mature form of matrix metalloproteinase 8 (mMMP8) would facilitate migration across matrix barriers *in vitro* and *in vivo*. We demonstrated that CAR T cells that co-express mMMP8 demonstrate robust migration across Matrigel and can kill tumor cells embedded in Matrigel *in vitro*. We tested the efficacy of these mMMP8 engineered cells in both leukemic and ovarian cancer cell models embedded in Matrigel in xenograft mouse models. Our results illustrate that unlike parental CAR T cells that have minimal anti-tumor efficacy in these models, CAR T cells that secrete mMMP8 promote T-cell infiltration, leading to the eradication of the tumors and survival. We anticipate that the co-expression of mMMP8 can be broadly utilized to improve the infiltration and efficacy of CAR T cells targeting many different antigens.

## Introduction

Chimeric antigen receptor T (CAR) cells are genetically modified autologous T cells that express chimeric receptors encoding an antigen-specific single-chain variable fragment (scFv), along with various costimulatory molecules^1,2^. Despite remarkable success in hematological cancer treatment, clinical experience with solid tumors has not been equally encouraging^3^. Several factors prevent the effectiveness of CAR T-cell therapy against solid tumors, including, but are not limited to, the (a) presence of an immunosuppressive environment, (b) heterogeneity in the expression of tumor-specific or tumor-associated antigens, (c) trafficking of immune cells to the tumor site, and (d) physical obstacles limiting T cell infiltration into the core of the tumor^4-6^.

One of the most fundamental barriers to the efficacy of immunotherapy is the physiochemical architecture of tumors. The extracellular matrix (ECM) is an integral part of the tumor stroma and consists of a variety of macromolecules, including glycosaminoglycans, proteoglycans, and fibrous proteins^7^. During cancer development, malignant cells contribute to ECM stiffness, and in return, the stiffened ECM alters the characteristics of cancer cells. This reciprocal relationship between the ECM and tumor cells ensures a hostile microenvironment that controls the supply of nutrients and oxygen, and most importantly functions as a barrier to the infiltration of immune cells. The tumor ECM presents a physical barrier restricting the entry of immune cells, such as T cells, into the tumor microenvironment, and the inability of T cells to infiltrate across the ECM has been reported in breast and lung cancers^8-10^. Since the infiltration of tumor-specific T cells into the tumor mass is a prerequisite for tumor killing and eradication, any barrier that prevents T cell infiltration renders ACT-based therapies ineffective, regardless of the functional capacity of the T cells. Recent studies using human T cells demonstrated that *ex vivo* expanded CAR T cells can display migratory exhaustion due to the downregulation of the ECM-degrading enzyme heparanase, and that overexpression of heparanase improved T cell infiltration and anti-tumor activity *in vivo*^11^. This study demonstrated the proof-of-concept T cells acquire phenotype of migratory exhaustion upon expansion. Unfortunately, heparin sulfate is only one component of the complex ECM in tumors, and enzymes with broader specificity for reversing migratory exhaustion are required.

Matrix metalloproteinases (MMPs) are an attractive group of ECM-degrading enzymes. MMPs have multifunctional roles, spanning from the digestion of ECM components, membrane shredding, chemokine processing, cancer, and even altering the activity of other proteases. Among the different MMPs, Matrix metalloproteinases 8 (MMP8) is attractive as a lead protease because it has a broad range of targets in the ECM, including aggrecan, gelatins, elastin, fibronectin, laminin, nidogen, and collagen^12^. Collagens are the most abundant proteins in the ECM^13,14^, they play a critical role in tumor cell growth, migration, and metastasis^8,15,16^. In this study, we tested the hypothesis that overexpression of the mature form of MMP8 can reverse the exhausted migratory phenotype of expanded CAR-T cells, leading to tumor infiltration and regression of solid tumors.

## Methods

### Cell lines

We cultured SKOV-3-CD19 and PlatGP cell lines in DMEM (Gibco, Invitrogen, Carlsbad, CA, USA) supplemented with 10% fetal bovine serum (FBS, HyClone, Thermo Scientific, Pittsburgh, PA, USA) and 2 mM GlutaMax (Invitrogen, Carlsbad, CA). Similarly, we cultured NALM6 cells obtained from ATCC in RPMI1640 (HyClone), supplemented with 10% FBS and 2 mM GlutaMax. All tumor cell lines were routinely tested for mycoplasma and surface expression of the target antigens.

### Microarray profiling

We extracted microarray profiling data from the Gene Expression Omnibus (GEO) under accession number GSE14352, which consisted of CD2^+^ T-lymphocytes from 10 healthy donors. RNA samples were extracted at 0, 24, and 72 h post-activation with CD3/CD28 Dynabeads. The Oligo package in R was used to extract raw expression values from the downloaded CEL files. To avoid batch variations between patients, we normalized the probe intensities using the Robust Multiarray Averaging (RMA) algorithm from the Oligo package, and plotted the normalized data using GraphPad Prism (v8.0). Statistical analysis was performed using the Wilcoxon rank-sum test.

### PBMC isolation and activation

We isolated peripheral blood mononuclear cells (PBMC) from anonymous buffy coats of healthy donors using Ficoll density separation and activated T lymphocytes with immobilized OKT3 (1 μg ml) and soluble anti-CD28 (0.5 μg ml) antibody (Abs) and then expanded them in complete medium containing RPMI1640 supplemented with 10% FBS. The cells were fed twice a week with recombinant interleukin-7 (IL-7) (10ng/mL) and interleukin-15 (IL15) (5ng/mL).

### Retroviral constructs, generation of retroviral producer cell lines and transduction of T lymphocytes

Mature MMP8 cDNA was cloned into a retroviral backbone along with GFP. We also cloned MMP8 into the CD19 CAR vector via E2A. The CD19 CAR vector contained CD8, 4-1BB, and CD3ζ endodomain. We transfected PlatGP cells stably expressing RD114 with retroviral constructs and enriched GFP-positive cells by flow cytometry to generate a stable retroviral producer cell line. The supernatant from these producer cell lines was used to transduce activated T-cells using retronectin-coated plates. We maintained the transduced T-cell lines in complete RPMI medium in a humidified atmosphere containing 5% CO2 at 37°C in the presence of IL-7 and IL15 for 2 weeks.

### T cell transwell migration assay

We assayed the ability of T cells to degrade and migrate across Matrigel using a transwell migration assay in a 96 well format (8 μm). We coated the wells with Matrigel and added 0.1M T cells that were activated for different time intervals to the upper chamber. The experiments were conducted in quadruplicate. After 24 h, we quantified the number of cells that migrated into the lower chamber using the MTT reagent.

For transwell migration assays performed in the presence of MMP8 or C/EBP inhibitors, we first activated T cells for 24 h with OKT3/anti-CD28 antibodies and simultaneously treated them with DMSO, MMP8 inhibitor, or C/EBP inhibitor. After 24 h, T cells were collected, added (0.1M/well) to the top chamber, and incubated for 24 h. The top plate was then removed, and the number of T cells that migrated into the lower chamber was quantified using MTT reagent.

### Bulk killing assay

We performed co-culture studies of targets and effectors to evaluate the in vitro efficacy of different CAR constructs to kill targets. FfLuc-NALM6 or FfLuc SKOV3-CD19 tumor cells were co-cultured with CAR T cells or MMP8 expressing CAR T cells at a 3:1 E:T ratio in 96 well. After 4 h, we added D-luciferin substrate to each well and measured luminescence with TopCount plate reader.

To perform a killing assay in Matrigel, 0.25×10^4^ FfLuc-NALM6 cells or 0.5×10^4^ FfLuc CD19-SKOV3-CD19 cells were seeded in 25μl Matrigel and allowed to solidify at 37°C °C for 30 min. Then, we laid 100μl of CAR T or MMP8 CAR T cells at a concentration of 0.75×10^6^ cells/ml on top of the Matrigel-enclosed target cells and incubated the plate for 18 h, and subsequently added D-luciferin substrate to quantify the live target cells with TopCount plate reader.

To simultaneously evaluate the migration and anti-tumor activity of CAR T cells, we coated Transwell inserts with Matrigel and laid 0.1 M CAR T cells/MMP8 CAR T cells on top of it and seeded target cells FfLuc-NALM6 cell (0.25×10^4^) or CD19 FfLuc SKOV3-CD19 (0.5×10^4^) in the lower chamber. The plate was incubated for 18 h, and D-luciferin substrate was added to the lower chamber to quantify the live target cells with TopCount plate reader. The experiments were performed in quadruplicates.

### RNA isolation and RT-PCR

To assess MMP8 transcript levels upon T cell activation, we isolated RNA from T cells that had been activated for different time intervals using OKT3/anti-CD28 antibodies, along with IL7 (10ng/mL) and IL15 (5ng/mL). CD14+ monocytes were used as positive control. Additionally, to assess *MMP8* transcript levels in T cells following treatment with a C/EBP inhibitor, T cells were activated and simultaneously treated with the inhibitor for 24 h.

Semi-quantitative RT-PCR was performed using 1000 ng of total RNA to prepare cDNA using the One Step PCR Master Mix Reagents Kit. We designed specific primers for *MMP8* and *GAPDH* (Glyceraldehyde-3-phosphate dehydrogenase), using 30 cycles for *MMP8* and 25 cycles for *GAPDH* PCR. The samples were loaded onto agarose gel to visualize the respective bands.

### Western blotting

To assay secreted MMP8, we harvested the supernatants from the cultures of T cells activated over different time intervals and confirmed the expression of MMP8 using western blotting using an MMP8 antibody. To assay MMP8 expression in T cells overexpressing MMP8, we harvested T cells transduced with GFP or MMP8 on day 10 and performed western blotting using anti MMP8 antibody and β-actin as the loading control.

### Collagen substrate degradation assay

We used the EnzyFluoTM Collagen Assay Kit (ECOL-100), according to the manufacturer’s protocol, to assay the activity of mMMP8 secreted by T cells. Briefly, 20 μL of 50 μg/mL collagen was incubated with 30 μL of supernatant media (diluted 10 times) from control T cells or MMP8 overexpressing cells cultured for 10 days. We then read the fluorescence at λex/em = 375/465 nm using Cytation.

### Flow cytometry

We performed flow cytometry analysis using the following Abs: CD3, CD4, CD8, CD45RA, CD45RO, and CD62L (all from Becton Dickinson, San Jose, CA, USA) conjugated with FITC, PE, PerCP, or APC fluorochromes. The samples were analyzed using a BD FACSCalibur.

### Mouse Model

All the mouse experiments were approved by the Institutional Animal Care and Use Committee of the University of Houston. We used the NSG mouse model to assess the *in vivo* anti-tumor effects of CAR and MMP8 CAR T cells. We subcutaneously injected either FfLuc-NALM6 or FfLuc-SKOV-3-CD19 (0.5 × 10^6^) resuspended in Matrigel into 10-week-old NSG mice. After 7 days of FfLuc-SKOV-3-CD19 injection and 30 days of FfLuc-NALM6 injection, CAR T cells were injected i.p. (5 × 10^6^ cells per mouse). We monitored tumor growth by bioluminescence *in vivo* imaging using the Xenogen-IVIS Imaging System. The investigator or veterinarian who monitored the mice three times a week detected signs of discomfort and euthanized the mice accordingly.

To investigate CAR T cell migration into tumors in mice, we injected FfLuc-SKOV-3-CD19 (0.5 × 106) resuspended in Matrigel into 10-week-old NSG mice. The tumor was allowed to grow for three weeks. Once the tumor was established, we injected the mice i.p. with 5 × 10^6^ CAR T cells (n=7) or MMP8-CD19-CAR T cells (n=7). Fourteen days after CAR T cell injection, we sacrificed the animals, harvested the tumors, and quantified the number of CAR T cells by flow cytometry using an antibody against human CD3.

### Statistical Analysis

Unless otherwise noted, we summarized the data as mean ± standard deviation (SD). To determine statistically significant differences between samples, we used a two-sided Student’s t-test with a p-value <0.05, indicating significance. For multiple comparisons, we used a repeated-measures analysis of variance (ANOVA) followed by a log-rank (Mantel-Cox) test. We analyzed the mouse survival data using the Kaplan-Meier survival curve to measure statistically significant differences. Before infusion of control or gene-modified T cells, we matched the mice based on the tumor signal for the control and treatment groups. To estimate the sample size, we considered the variation and mean of the samples to use the smallest possible sample size. We estimated the sample size to detect a difference in means of two standard deviations at a significance level of 0.05, with 80% power. Prism version 9 software (GraphPad, La Jolla, CA, USA) was used for graph generation and statistical analyses.

## Results

### Ex vivo expanded T cells showed migratory exhaustion

To quantify the correlation between T-cell activation and migratory phenotype, we first compared the ability of T cells, activated, and *ex vivo* expanded over different intervals of time, in their ability to degrade Matrigel using a transwell-based migration assay. Matrigel, a basement membrane matrix extracted from Engelbreth–Holm–Swarm sarcomas, is predominantly composed of three major ECM proteins, laminin (60%), collagen IV (30%), and entactin (8%)^17^. T cells activated briefly for 24 h showed robust migration across Matrigel, which diminished as they expanded beyond 48 h (**Fig 1A**). This data is consistent with previous studies that demonstrated that long-term *ex vivo* expansion attenuates the ECM degradation capabilities of activated human and rodent T cells^12^. Among the various human MMPs, we investigated MMP8 for two reasons: (1) unlike most members of the MMP family that are pro-tumorigenic, several studies have shown that MMP8 is predominantly anti-tumorigenic, and (2) MMP8 is expressed in immune cells ^18-20^. We performed RT-PCR on human T cells harvested at different time points after activation and observed that T cells activated beyond 48 h showed reduced *MMP8* transcript levels (**Fig 1B**). We validated these results at the protein level by determining secreted MMP8 protein in the same cell populations and observed an initial spike in MMP8, which decreased with time (**Fig 1C**). To confirm the causative role of MMP8 in promoting T cell migration, we performed a transwell migration assay across Matrigel with or without the MMP8 specific inhibitor, MMP8-Inhibitor-I (no known activity against other MMPs observed in vitro)^21^. MMP8 inhibition reduced the migratory capacity of T-cells (**Fig 1D**).

**Figure 1.**
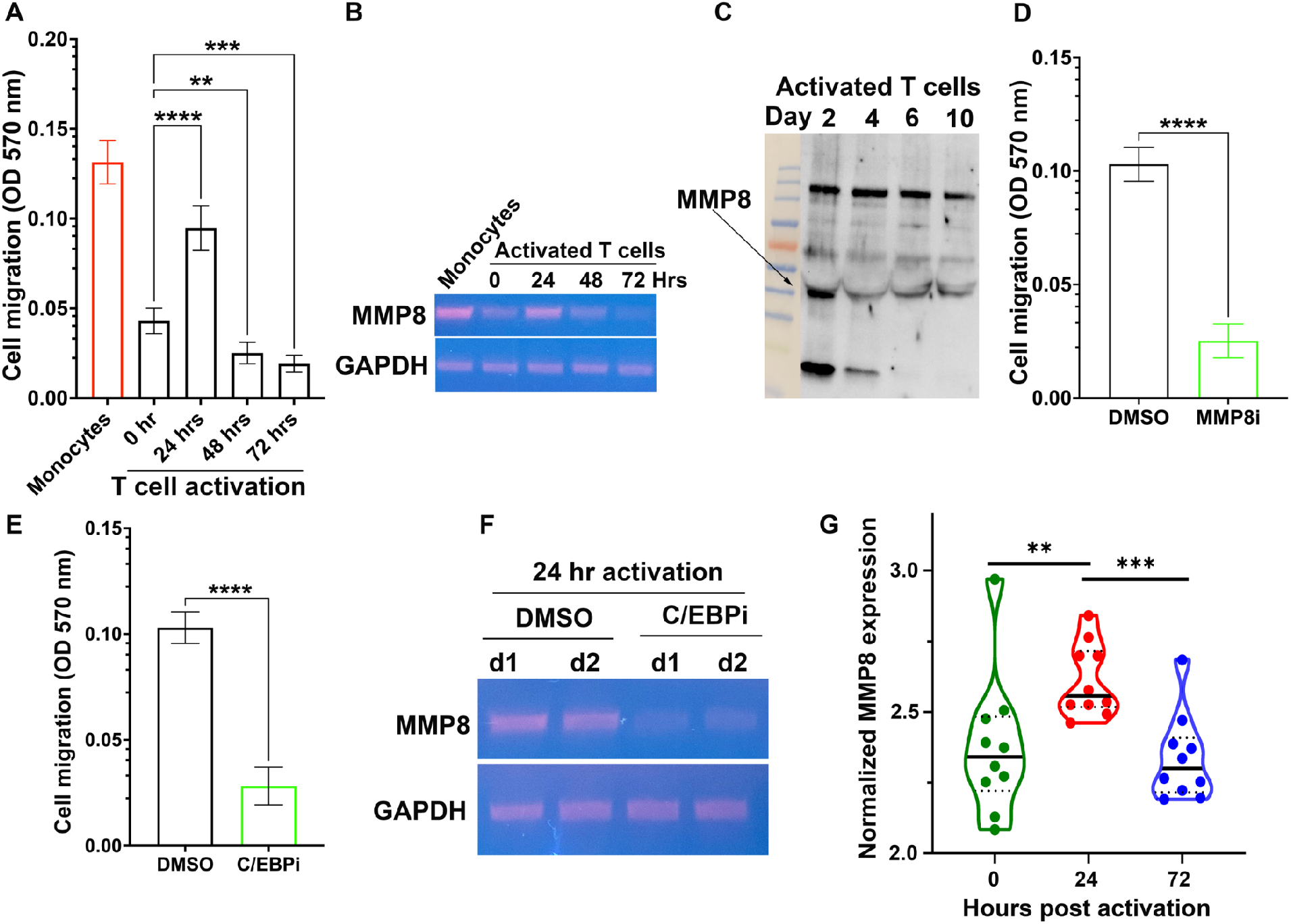
*Ex vivo* expansion of T cells induced migratory exhaustion and downregulated MMP8. **(A)** The migration of monocytes, freshly isolated resting T cells, and activated T cells across Matrigel over 24h evaluated using a standard transwell assay. **(B)** RT-PCR of *MMP8* expression within T cells after from activation and expansion. **(C)** Western blot of T cell supernatants testing for MMP8 **(D-E)** The impact of (D) MMP8 inhibitor I and (E) C/EBP inhibitor on the migration of activated T cells across Matrigel over 24 hours evaluated using a standard transwell assay. The data is from two separate donors. **(F)** RT-PCR of *MMP8* expression within T cells upon C/EBP inhibition. **(G)** Violin plot showing normalized *MMP8* expression (n = 10 donors) at 0, 24, and 72 hours post T cell activation via microarray profiling. The bold line represents the median and the dotted lines represent the upper and lower quartiles. Statistical testing was performed using the Wilcoxon rank sum test. *p < 0.05, **p < 0.01. ***p < 0.001, ****p < 0.0001.

Since the RT-PCR data showed that the transcripts of *MMP8* were downregulated upon activation (**Fig 1B**), we wanted to investigate the impact of transcription factors (TF) that can regulate MMP8. The promoter region for MMP8 contains a recognition site for the master TF, CCAAT/enhancer-binding protein beta, C/EBP-β^22^. To investigate the role of C/EBP-β in regulating MMP8 expression, we treated activated T cells with C/EBP-β inhibitor and tested alterations in their migratory capacity and MMP8 expression. C/EBP-β inhibition significantly reduced *MMP8* transcript levels and the migratory ability of T cells (**Fig 1E-F)**.

To generalize our results across T cells derived from a panel of human donors, we used microarray data from a publicly available cohort (GSE14352)^23^. We extracted *MMP8* expression data from 10 donors at different stages of activation **(Supplemental Fig 1)**. Consistent with our experimental data, the microarray data showed a significant upregulation of *MMP8* at 24 h post-activation compared with baseline expression (**Fig 1G**). Furthermore, *MMP8* expression declined between 24 and 72 hours post-activation. Taken together, these data show that although MMP8 expression is increased upon activation, this effect is transient, and further *ex vivo* expansion reduced MMP8 expression and leads to migratory exhaustion of T cells.

### Overexpression of MMP8 rescued migratory exhaustion of *ex vivo* expanded T cells

Since our data suggest that *ex vivo* expansion of T cells reduced MMP8 secretion and migration of T cells, we hypothesized that overexpression of mature MMP8 (mMMP8) could rescue the migratory exhaustion of *ex vivo* expanded T cells. Accordingly, we retrovirally transduced T-cells with either GFP or mMMP8-E2A-GFP. We confirmed the expression of GFP and inferred the expression of MMP8 in T cells using flow cytometry (**Fig 2A**). To assay secreted MMP8, we harvested the supernatants from the culture of T cells activated over different intervals of time and confirmed the expression of MMP8 using western blotting with an MMP8 antibody (**Fig 2B**). To assay the enzymatic activity of secreted mMMP8, we used a fluorometric collagenase assay that utilizes a quenched collagen substrate that releases quantifiable fluorescence upon proteolysis. T cells transduced with mMMP8 showed significantly increased collagen degradation compared with T cells transduced with GFP (**Fig 2C**). To determine whether overexpression of mMMP8 could rescue migratory exhaustion, we *ex vivo* expanded both sets of T cells (GFP and MMP8-E2A-GFP) for 10 days. When assayed using the standard Transwell migration assay, mMMP8 T cells showed significantly improved migration compared to GFP T cells (**Fig 2D**). We again confirmed that this migration could be attributed to the overexpression of MMP8 using MMP8-inhibitor-I (**Fig 2E**). Collectively, our data illustrate that the overexpression of mMMP8 in T cells restores migration.

**Figure 2.**
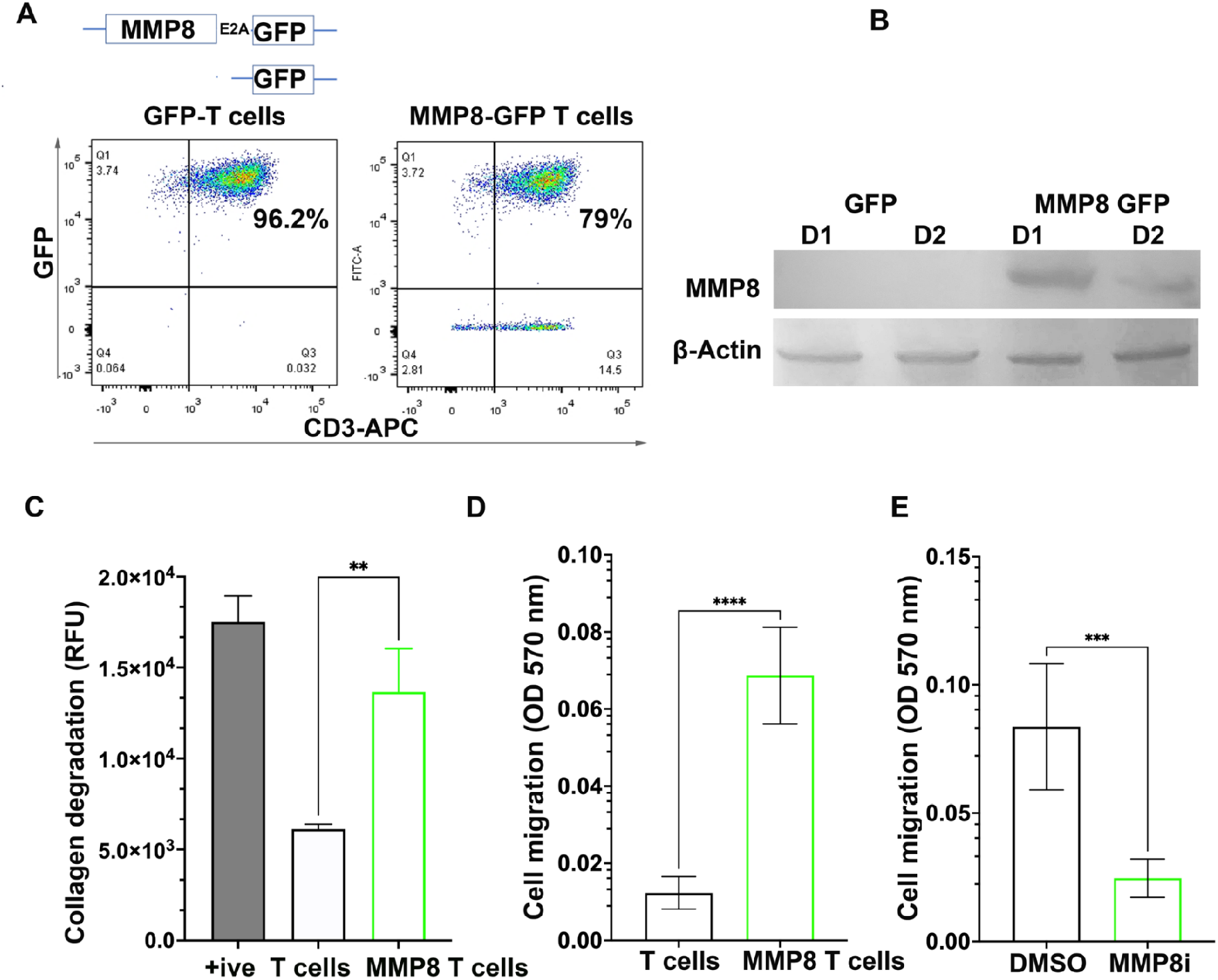
T cells overexpressing MMP8 showed superior infiltration compared to T cells alone *in vitro*. **(A)** Schematics of MMP8 construct along with flow cytometric quantification of MMP8 expression (GFP) in T cells. **(B)** Western blot of T-cell lysates from two separate donors harvested at day xx after expansion. **(C)** Collagen degradation mediated by T cells measured by EnzyFluoTM Collagen Assay Kit. **(D)** The migration of T cells and MMP8 OE T cells across Matrigel over 24h evaluated using a standard transwell assay. We quantified the number of cells that migrated to the lower chamber by MTT reagent. **(E)** The impact of MMP8 inhibitor I on the migration of activated T cells across Matrigel over 24 hours evaluated using a standard transwell assay. All data in Figure 2 was obtained from at least two different donor-derived T-cell populations.

### Overexpression of MMP8 rescued migration and killing ability of *ex vivo* expanded CAR T cells

Having established that mMMP8 OE can facilitate T cell migration, we next investigated the impact of mMMP8 expression in *ex vivo* expanded tumor-targeting T cells. Accordingly, we generated CD19R-4-1BB-CD3z (CD19-CAR) and MMP8-E2A-CD19R-4-1BB-CD3z (MMP8-CD19-CAR) retroviral constructs (**Fig 3A**). We transduced activated T cells to generate CD19-CAR and MMP8-CD19-CAR T cells, which were expanded over a 10-day period. We performed phenotyping of both populations of CAR T cells by flow cytometry and confirmed that MMP8 overexpression did not alter the memory phenotype of CD19-CAR T cells (**Fig 3B**). Similarly, there were no differences between the proliferation of CD19-CAR T cells and MMP8-CD19-CAR T cells (**Fig 3C**).

**Figure 3.**
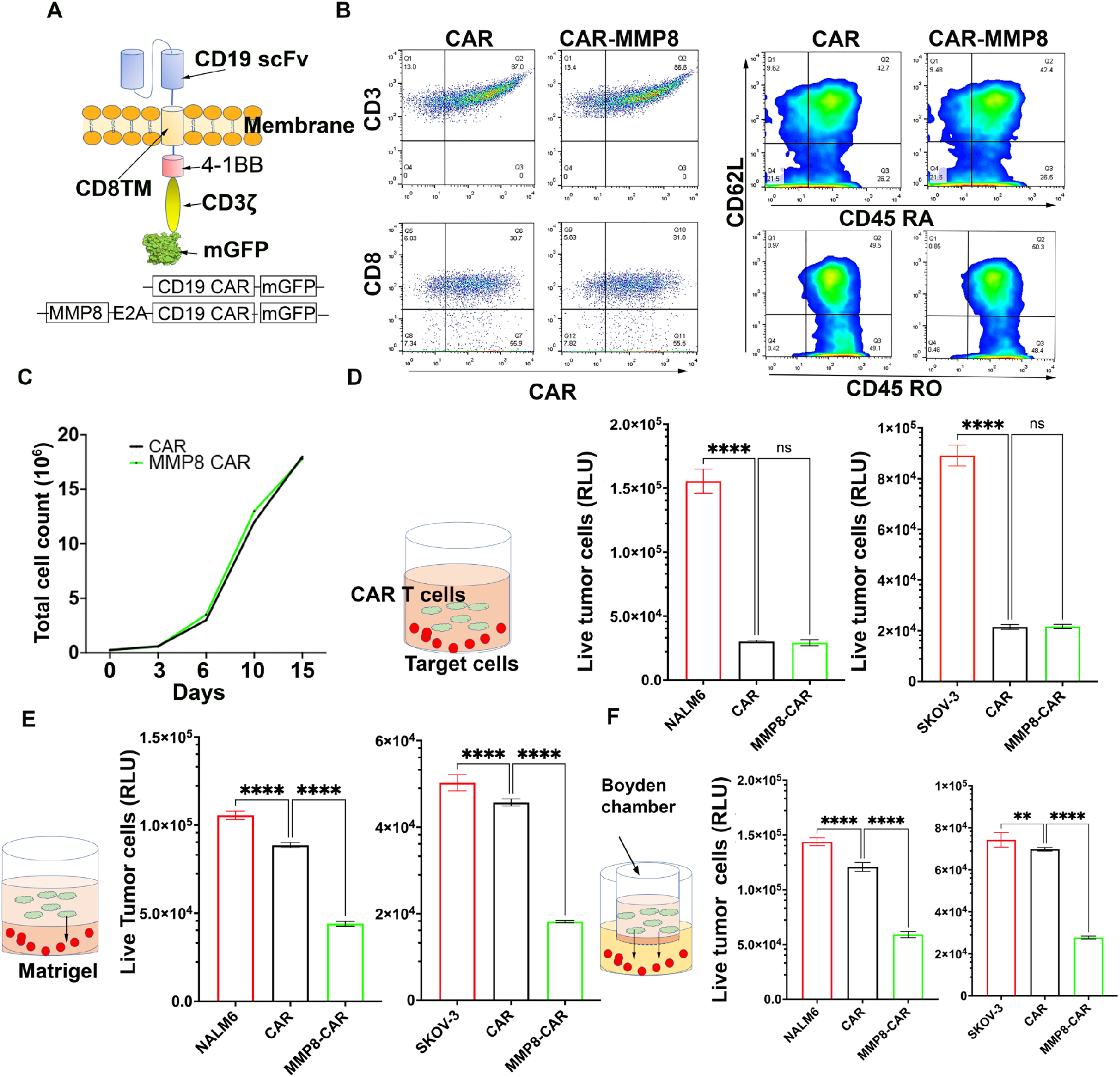
CD19 CAR T cells T co-expressing *MMP8* showed superior infiltration and tumor killing efficacy compared to CAR T cells alone *in vitro*. **(A)** Schematics of CD19 CAR and MMP8 CD19 CAR retroviral constructs. We fused GFP to the CD3ζ terminal of CAR to facilitate monitoring of transduction. **(B)** Phenotyping of CD19 and MMP8 CD19 CAR T cells by flow cytometry to evaluate the expression of CAR and memory phenotypic markers: naive (CD45RA+), central-memory (CD45RO+CD62L+) and effector-memory (CD45RO+CD62L−) T cells. **(C)** Impact of MMP8 on cell proliferation of CAR T cells. We cultured MMP8-CD19 and CD19 CAR T cells for 15 days and counted the cells at different time points. **(D)** Impact of MMP8 on killing efficacy of CAR T cells of targets. Coculture of CD19 CAR/MMP8-CD19 CAR with targets (FfLuc-SKOV3-CD19/FfLuc-NALM6) at 3:1 effector:target (E:T) ratio for 4 hours. We quantified live targets using D-Luciferin substrate with TopCount scintillation counter. **(E)** Impact of MMP8 on killing efficacy of CAR T cells of targets in the presence of Matrigel. We seeded FfLuc expressing targets in Matrigel in 96-well plate and laid MMP8-CD19CAR or CD19 CAR T cells on top of them at 3:1 E:T ratio. After 18 hours of incubation, we added D-Luciferin substrate to the cells to quantify luminescence, indicative of live cells, with TopCount scintillation counter. **(F)** Simultaneous evaluation of CAR T cell migration and killing of targets using transwell assay. We seeded MMP8-CD19 CAR or CD19 CAR T cells in the upper chamber coated with Matrigel and the targets in the lower chamber at 3:1 E:T ratio. After 18 hours of incubation, we added D-Luciferin substrate to cells in the lower chamber to quantify luminescence, indicative of live cells, with TopCount scintillation counter. All the data in Figure 3 was obtained from at least two different donor-derived T cell populations.

To assay the anti-tumor killing functionality of CAR T cells *in vitro* we set up a panel of three separate experiments. First, we performed a conventional killing assay by co-culturing CD19-CAR T cells or MMP8-CD19-CAR T cells with either FfLuc-NALM6 (leukemia) or FfLuc-SKOV-3-CD19 (ovarian) tumor cells (**Fig 3D and Supplemental Figure 2**). As expected, both sets of CAR T cells showed significant killing of both tumor targets, and their killing rates were comparable (**Fig 3D**). Second, to model the impact of ECM on tumor cell protection, we pre-encapsulated either FfLuc-NALM6 or FfLuc-SKOV-3-CD19 tumor cells in Matrigel and then layered CAR T cells on top (**Fig 3E**). As expected, embedding the tumor cells in Matrigel decreased their susceptibility to killing by CAR-T cells. Under these conditions, MMP8-CD19-CAR T cells showed significantly better killing than CD19-CAR T cells, likely due to their ability to degrade Matrigel via MMP8 secretion (**Fig 3E**). Third, we set up a killing assay in which CAR T cells were embedded in Matrigel and seeded in the top well of a transwell chamber (**Fig 3F**). Tumor cells (FfLuc-NALM6 or FfLuc-SKOV-3-CD19) were seeded in the bottom well. In this setup, CAR T cells have to degrade the Matrigel and migrate across the pores of the Transwell insert before killing the tumor cells. Not surprisingly, MMP8-CD19-CAR T cells were superior to CD19-CAR T cells in killing tumor cells (**Fig 3F**). Taken together, these results illustrate that MMP8-CD19-CAR T cells show a similar frequency of memory and proliferative cells to CD19-CAR T cells, but show enhanced migratory capacity in being able to degrade the ECM to enable the killing of tumor cells.

### MMP8-CD19 CAR T cells showed potent tumor infiltration and anti-tumor efficacy in mouse tumor models

Since the in vitro killing assays illustrated the role of MMP8 in the migration of T cells, leading to tumor cell killing, we next evaluated the anti-tumor efficacy of CAR T cells against a solid tumor model *in vivo*. We subcutaneously injected 0.2 × 10^6^ FfLuc-SKOV-3-CD19 tumor cells in Matrigel into NSG mice. Once the tumor became palpable (5-7 days), we randomly divided the mice into three groups based on IVIS imaging. One group, the tumor alone group (n = 5), was left untreated as a control. The second (n = 7) and third (n = 8) groups received 5 × 10^6^ CD19-CAR T cells and MMP8-CD19-CAR T cells per mouse, respectively. Tracking the tumor burden as a function of time showed that the parental CD19-CAR T cells showed very little activity against the FfLuc-SKOV-3-CD19 tumor (**Fig 4A**), and by day 25, all animals in both the tumor-only group and the tumors treated with CD19-CAR T cells died (**Fig 4B**). In contrast, on day 28, all animals with FfLuc-SKOV-3-CD19 tumors treated with MMP8-CD19-CAR T cells were completely tumor-free, and 100% of the animals survived to day 50 when the experiment was terminated (**Fig 4C**).

**Figure 4.**
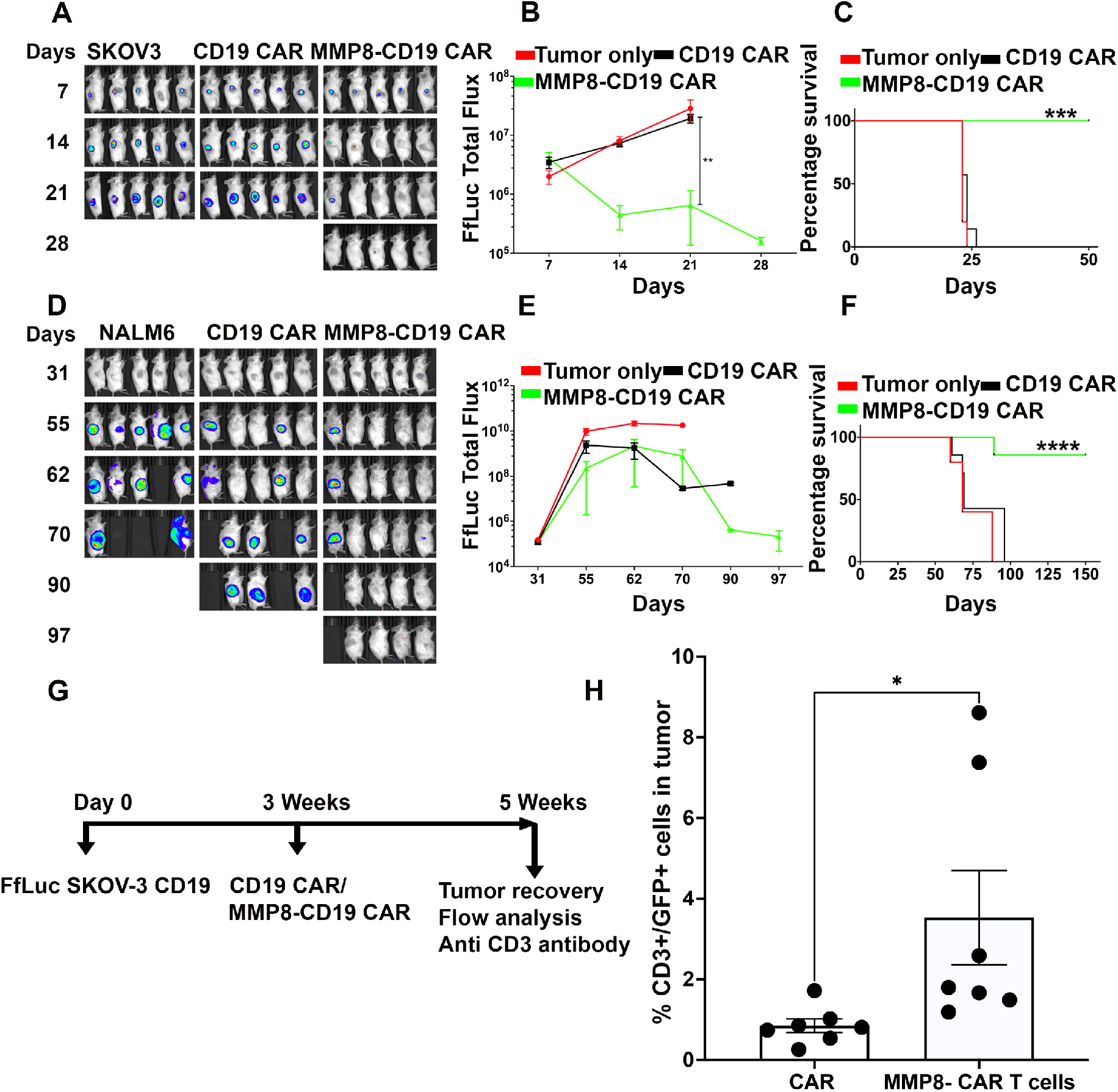
MMP8-CAR T cells showed superior infiltration and improve overall survival in xenograft tumor models. (**A-C**) Tumor flux imagine (A) and survival (B) analysis of mice engrafted subcutaneously with FfLuc SKOV-3 CD19 and treated i.p. with of either CD19 CAR or MMP8 CD19 CAR T cells.; control n=5, CD19 CAR n=7, MMP8 CD19 CAR n=8 mice **(D-F)** Tumor flux imaging and survival analysis of mice engrafted subcutaneously with FfLuc NALM6 and treated i.p. with either CD19 CAR or MMP8 CD19 CAR T cells.; control n=5, CD19 CAR n=7, MMP8 CD19 CAR n=7 mice **(G-H)** Flow analysis of CD3+ T and GFP+ T cells detected within the FfLuc-SKOV3-CD19 mice tumors (G) schematics (H) histogram of % CAR T cells within tumor summarize means ± SEM of 7 mice each from CAR T cell alone or MMP8 CAR T cell treated mice. The data shown is from two independent repeats.

Next, we evaluated the anti-tumor efficacy of CAR T cells against NALM6 tumors in NSG mice. We subcutaneously injected 0.2 × 10^6^ FfLuc-NALM6 tumor cells into Matrigel in mice. On day 7, mice were randomly divided into three groups based on IVIS imaging. Unlike the FfLuc-SKOV-3-CD19 tumor cells described above, the NALM6 cells consistent with their leukemic origin when embedded in Matrigel, yielding slow-growing tumors (**Fig 4D**). CD19 CAR T cells had a transient anti-tumor effect, but this did not yield an improvement in survival compared to the mice that only received the tumor (**Fig 4E**). Treatment of tumor bearing mice with MMP8-CD19-CAR T cells yielded substantial anti-tumor activity that was durable, and 7/8 mice survived to day 150 when the experiment was terminated (**Fig 4F**).

Our *in vitro* data showed no differences in the memory phenotype or proliferative capacity between CD19-CAR T cells and MMP8-CD19-CAR T cells. To investigate whether the anti-tumor efficacy of MMP8-CD19-CAR T cells was due to the enhanced migratory potential of these CAR T cells, leading to better tumor infiltration, we established FfLuc-SKOV-3-CD19 tumors in Matrigel in NSG mice for 21 days (longer duration led to large tumors). We subsequently injected MMP8-CD19-CAR T cells or CD19-CAR T cells into mice and fourteen days after CAR T cell injection, the animals were sacrificed, tumors were harvested, and the frequency of CAR T cells was quantified using an antibody against human CD3 (**Fig 4G, Supplemental Fig 3**). As expected, on average, 3.9-fold more MMP8-CD19-CAR T cells were recovered from the tumor than CD19-CAR T cells (**Fig 4H**). Collectively, these results establish that OE of mMMP8 promotes the degradation of Matrigel, facilitating the migration of CAR T cells into the tumor, yielding durable anti-tumor efficacy.

## Discussion

Despite success with hematological cancers, T cell-based therapies encounter significant obstacles in treating patients with solid tumors. During cancer progression, tumor cells foster the development of an immunosuppressive tumor microenvironment (TME), working in concert with fibroblasts, pro-tumor macrophages, and endothelial cells embedded within a robust extracellular matrix (ECM)^24,25^. The ECM is a dominant contributor to tissue architecture and can function as a physical barrier to prevent the entry of anti-tumor T cells, facilitating escape from immunosurveillance^25,26^. The ECM is composed of collagens, proteoglycans, fibronectin, and glycosaminoglycans and is a highly dynamic structure that is continuously remodeled by the cells of the TME to sustain tumor growth while sustaining immunosuppression.

The physiochemical properties of the ECM in the TME differ from those of the ECM in normal epithelial tissues. In normal tissue, the collagen matrix exists in a relaxed state and provides a guiding framework for T-cell migration and immunosurveillance. In contrast, high interstitial pressure and high expression of collagen-processing lysyl oxidases and MMPs increase collagen deposition and remodeling of ECM fibers. In addition to collagen, the deposition of other ECM components, including hyaluronan, fibronectin, and tenascin C, results in desmoplasia, a highly fibrotic phenotype that mimics organ fibrosis. Desmoplasia and ECM remodeling including excess production of hyaluronan and collagen, are characteristics of cancers such as breast and pancreatic cancers associated with poor prognosis ^10,27^. In breast cancer patients, there is a positive correlation between tissue stiffness and malignant phenotypes, including tumor size, histologic grade, and estrogen receptor (ER) status, with triple-negative breast cancer tissues ranking the stiffest^28^. In pancreatic ductal adenocarcinoma, there is a strong negative correlation between patient survival and dense structure of ECM^29^. From a therapeutic perspective, T-cell therapies that aim to mediate tumor regression must be able to navigate through this ECM barrier to execute their anti-tumor functions.

MMPs are a class of hydrolytic enzymes that naturally degrade collagen and other components of the ECM and play an active role in the remodeling, leading to tumor invasion and metastasis^30,31^. Similarly, immune cells, specific cells of the monocytic lineage, and neutrophils secrete MMPs to facilitate infiltration. Activated T cells upregulate the expression of MMP transcripts and, at least *in vitro*, the upregulation of MMP transcripts is associated with efficient migration of cytotoxic T lymphocytes (CTL) through a collagen layer in transwell assays, and this activity could be blocked with the aid of a pan-MMP inhibitor^32^. *In vivo*, degradation of collagen can serve additional anti-tumor functions since the collagen fragments arising from proteolytic cleavage can activate integrin-dependent T cell motility and establish a chemotactic gradient that guides T cells towards tumor cells^32^. Despite this promise, as our studies illustrate, T-cell activation only transiently upregulates MMP expression and is rapidly lost even 48h after activation, leading to T cells acquiring migratory exhaustion upon expansion for cell therapy manufacture.

To design T cells to secrete MMPs, we factored into both the pro- and anti-tumorigenic properties of these proteolytic enzymes. Most MMPs are pro-tumorigenic and can serve as biomarkers for advanced cancers^18,19^. Cancer studies have shown that *MMP1, MMP11*, and *MMP13* are elevated in several different types of cancers, whereas *MMP8* is rarely detected^33^. In animal models, *MMP8*^*−/−*^ mice have a high incidence of skin tumors, which can be rescued by bone marrow transplantation. This study clearly demonstrated the anti-tumor role of MMP8 *in vivo*^34^.

Mechanistically, the protective function of MMP8 was likely due to the enzymatic processing of mediators of inflammation rather than its known role in degrading collagen fibrils, emphasizing the multiple positive effects of MMP8 activity^34,35^. Further evidence for the anti-tumor properties of MMP8 arises from genetic manipulation of MMP8 in human breast cancer cells, wherein overexpression of MMP8 in metastatic cell lines reduced the metastatic potential of cells, while its downregulation had the opposite effect on non-metastatic cells^36^. Similar findings in syngeneic melanoma and lung carcinoma have established the role of MMP8 as a promoter of metastatic suppression via the modulation of cell adhesion and invasion^37^. Collectively, most studies support an anti-tumor role for MMP8, and since immune cells are known to express MMP8, we chose it as the enzyme of choice for the degradation of ECM, facilitating T-cell infiltration.

Our experimental results clearly validated our hypothesis that overexpression of the mature, secreted form of MMP8 enabled T-cell and CAR T-cell migration across Matrigel *in vitro. In vivo*, MMP8-CD19 CAR T cells showed robust infiltration of solid tumors in xenograft mouse models, leading to potent anti-tumor efficacy and tumor-free survival. We recognize that combating solid tumors is a multifactorial problem, and migration leading to infiltration solves only one of the immunosuppressive barriers that T-cell therapies encounter^38,39^. However, migration leading to infiltration is the most fundamental barrier, since T cells excluded from tumors have no opportunity to execute any anti-tumor functions. By solving the barrier of infiltration, we provide T cells with the opportunity to interact with tumor cells. While other immunosuppressive factors can still facilitate T-cell exhaustion, the presence of T cells within the tumor allows us to deploy other therapeutics in our arsenal, including immune checkpoint antibodies, to facilitate an anti-tumor response^38,39^. We anticipate that genetic modifications to express mMMP8 can serve as a fundamental module to facilitate the next generation of T and potentially NK cell therapies^40^.

## AUTHORSHIP CONTRIBUTION

Conceived and designed the study: IB, NV

Prepared the manuscript: IB, HS, NV

Performed experiments: IB

Analyzed data: IB, MJM, NV

All authors edited and approved the manuscript.

## ACKNOWLEDGEMENTS

This study was supported by the National Institute of Health (R01GM143243). We would like to acknowledge the MDACC Flow Cytometry and Cellular Imaging Core facility for FACS sorting (NCI P30CA16672), Intel for the loan of computing cluster, and the UH Center for Advanced Computing and Data Systems (CACDS) for high-performance computing facilities.

## Supplemental Figures

**Supplemental Figure 1.**
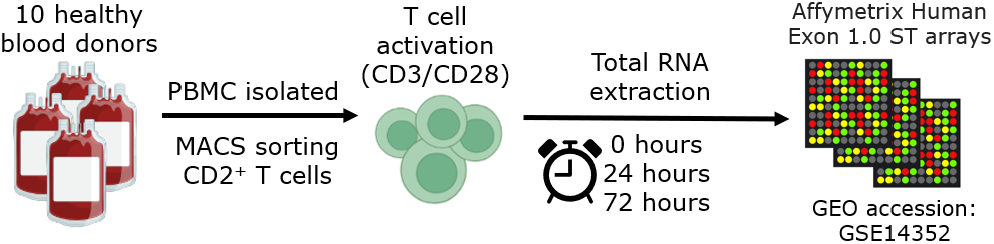
The GSE14352 dataset is derived from PBMCs isolated from ten healthy donors were activated using CD3/CD28 for 0, 24, and 48 h. The RNA was extracted at each time point, converted to labeled cDNA, and hybridized to Affymetrix Human Exon 1.0 ST arrays.

**Supplemental Figure 2.**
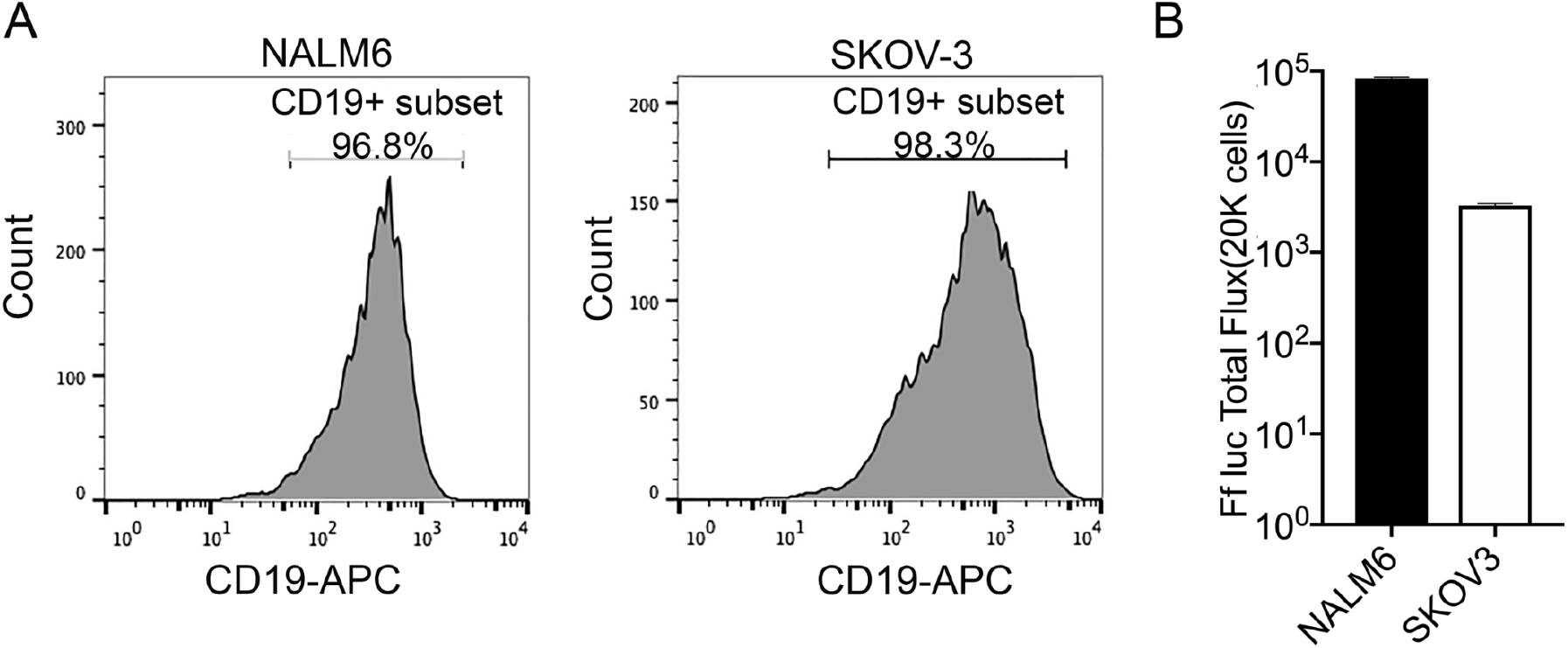
Evaluation of CD19 surface marker and Ffluc expression in targets. **(A)** Phenotyping of FfLuc-NALM6 and FfLuc-SKOV3-CD19 target cells was performed by flow cytometry using fluorescent antibody against surface CD19. More than 95% of cells expressed surface CD19. **(B)** NALM6 and SKOV-3-CD19 cells were engineered to express GFP FfLuc and assayed using D-luciferin substrate.

**Supplemental Figure 3.**
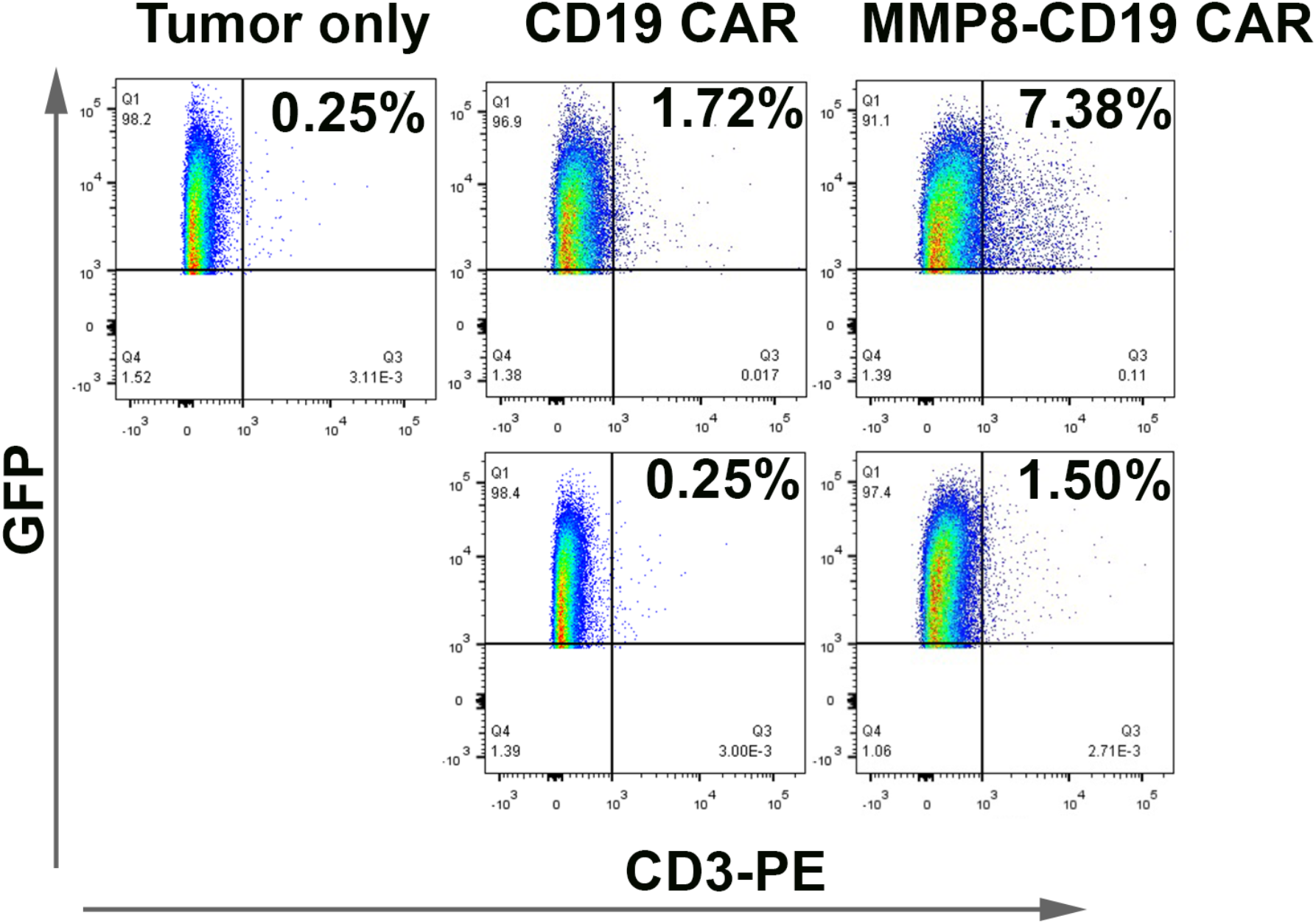
Flow analysis of CD3+ and GFP+ CAR T cells detected within the FfLuc-SKOV3-CD19 mice tumors.

